# Genome assembly and annotation of the naked mole rat *Heterocephalus glaber* reared in Japan

**DOI:** 10.1101/2025.05.20.654782

**Authors:** Kouhei Toga, Kaori Oka, Hiroyuki Tanaka, Takehiko Itoh, Atsushi Toyoda, Hidemasa Bono, Kyoko Miura

## Abstract

The naked mole rat (NMR, *Heterocephalus glaber*) is a eusocial rodent that is native to northeastern Africa. NMRs exhibit extraordinary traits such as longevity, resistance to age-related decline, and remarkable hypoxia tolerance. Although the reference genome of this species has been determined because of its unique characteristics, the significance or role of intraspecific genomic variations remains unknown. In this study, we used PacBio long-read sequencing to generate a genome assembly of NMR reared in Japan. The assembled genome is 2.56 Gb. Benchmarking Universal Single–Copy Orthologs (BUSCO) revealed high completeness (95.2%). BRAKER3 estimated 26,714 protein-coding genes, and we successfully added functional annotations for 26,232 protein-coding genes using the functional annotation workflow. We identified 417 gene models that were previously undetectable in the reference genome of this species. We also identified structural and amino acid sequence variations between our assembly and the reference genome, suggesting the presence of intraspecific genomic variations. This new genomic resource could help uncover the molecular mechanisms underlying the behavioral and physiological traits of NMR.

## Background & Summary

The naked mole rat (NMR) has received significant attention in the field of biology because of its unique behavior and physiology. NMR is the longest-lived rodent, exhibiting extraordinary longevity and resistance to age-related diseases, including cancer, with a maximum lifespan of 40 years despite its small size^1^. This remarkably long lifespan deviates from the typical positive correlation observed between body mass and lifespan in mammals^2^. The NMR is a eusocial rodent that lives cooperatively in subterranean colonies consisting of a single breeding female, one to three breeding males, and non-reproductive workers^3^. Adapted to group-living in semi-enclosed subterranean environments, they exhibit significant tolerance to both hypoxia and anoxia^4^. Comparative genomic analysis between NMR and other mammals revealed that genomic mutations in NMR contribute to longevity, adaptation to the subterranean environment, and resistance to carcinogenesis^5–9^. Additionally, amino acid substitutions are enriched at disease-causing sites in human^6^. Therefore, the NMR genome is an important resource for identifying genetic factors related to its unique ecology, physiology, and resistance to age-related diseases.

The concept of a reference genome is based on a single representative genome sequence for each species and is broadly applied to many organisms, as seen in the Earth BioGenome Project^10^. However, this concept does not account for intraspecific variation. Therefore, it remains unclear how genomic structures are preserved within the same species. Supporting these limitations, intraspecific genomic variations and chromosomal rearrangements have been observed in mammals, larvaceans, and insects^11–13^. Notably, maximal genetic distances within NMR species are approaching interspecific values observed among other Bathyergid species, based on calculations for the cytochrome-b gene between the Ethiopian and southern Kenyan populations^14^. This suggests the accumulation of genetic differences within the NMR species. Additionally, the rivers in Kenya may cause reproductive isolation, leading to local genetic differences^15^. However, genome resources are currently insufficient to identify genomic differences in NMR species. To address this gap, we sequenced the whole genome of a single NMR individual reared in Japan using Illumina short reads and PacBio long reads. Gene prediction and functional annotation were performed using BRAKER3^16^ and the functional annotation workflow (Fanflow)^17^ pipeline. We verified the presence of variations at both the genome and gene levels between NMR species, demonstrating that our genome assembly provides a valuable resource for advancing the understanding of genomic functions related to NMR physiology and behavior.

## Methods

### Sample collection, sequencing, and genome assembly

NMRs were maintained at 30 ± 0.5 °C and 55 ± 5% humidity with 12 h light and 12 h dark cycles at Kumamoto University. All animal experiments were performed in accordance with the Guide for the Care and Use of Laboratory Animals (United States National Institutes of Health). The ethics committee of Kumamoto University approved all the procedures (approval No. A2020-042, A2022-079 and A2024-063). Genomic DNA was extracted from the quadriceps muscle tissues using a QIAGEN Genomic-tip (QIAGEN, Hilden, Germany) according to the manufacturer’s instructions. The quality and quantity of the extracted genomic DNA were assessed using a Qubit 4 fluorometer (Thermo Fisher Scientific, Waltham, MA, USA) and a Pippin Pulse system (Agilent Technologies). DNA was sheared into 30–100 kb fragments using a g-tube device (Covaris Inc., MA, USA). A continuous long-read (CLR) single-molecule real-time (SMRT) bell library was prepared using the SMRTbell Express Template Prep Kit 2.0 (Pacific Bioscience, Menlo, CA, USA) according to the manufacturer’s instructions. The CLR libraries were size-selected using the BluePippin system (Saga Science, MA, USA) with a lower cutoff of 30 kb. The library was run on the PacBio Sequel II platform using two SMRT Cell 8Ms with Binding Kit 2.0, and Sequencing Kit 2.0 with 20-h movies. Additionally, genomic DNA was fragmented to an average size of 500–600 bp using a focused ultrasonicator M220 (Covaris Inc., MA., USA). Paired-end libraries with 450–550 bp insert sizes were constructed using the TruSeq DNA PCR-Free Library Prep Kit (Illumina, CA, USA) and size-selected on an agarose gel using a Zymoclean Large Fragment DNA Recovery Kit (Zymo Research, CA., USA). The final libraries were sequenced following the 2 × 250 bp paired-end protocol for NovaSeq 6000 systems (Illumina, San Diego, CA, USA).

Canu 2.1^19^ was used for *de novo* assembly of PacBio CLR reads with parameters ‘-genomesize 3.0g - coverage 200 -minlength 1000 -raw -pacbio’. Three rounds of polishing were performed using Pilon^20^ with the aforementioned Illumina short reads. Purge_dups^13^ was used with default parameters on the polished assembly to purge haplotigs.

### Gene prediction and functional annotation

Since repetitive sequences can lead to overestimation of gene predictions, repetitive sequences were filtered using RepeatModeler v2.0.3 and RepeatMasker v4.0.6 in Extensive *de novo* TE Annotator v2.0.1^23^, and were soft-masked. We identified 978 Mb of repetitive sequences, accounting for 38.19% of the genome assembly^24^. Gene prediction was performed using BRAKER3^16^. Public RNA sequencing (RNA-Seq) data^25^ and Vertebrate protein dataset (https://bioinf.uni-greifswald.de/bioinf/partitioned_odb11/, accessed on October 10, 2023) were used as extrinsic evidence for BRAKER3^16^. Metadata of public RNA-Seq data was collected using SRA Run Selector in National Center for Biotechnology Information (NCBI), and manually filtered to retain only samples with well-defined metadata. Sequence read archive (SRA) retrieval of public RNA-Seq and conversion to FASTQ files was performed using SRA toolkit v3.0.5 (https://github.com/ncbi/sra-tools, accessed on June 30, 2023). Trimming and quality control of the FASTQ files were performed using Trim Galore! v.0.6.10^26^ with ‘--fastqc, --trim1’ option. Mapping RNA-Seq reads to the genome was performed using HISAT2 v2.2.1^27^ with ‘-q, --dta’ option, and then obtained SAM files were converted to BAM files using SAMtools v1.12^28^. The obtained BAM files were supplied to the BRAKER3 workflow. Gene set coverage was assessed using BUSCO v5.2.2^22^ with ‘-l glires_odb10.’ Fanflow^17^ was used as the reference protein set in Ensembl release-111. This workflow used GGSEARCH v36.3.8g in the FASTA package (https://fasta.bioch.virginia.edu/, accessed on July 19, 2023) to compare amino acid sequences against datasets from humans, mice, guinea pigs, female NMR, male NMR, and UniProtKB/Swiss-Prot. Domain searches were conducted using HMMSCAN in HMMER v3.3.2, with Pfam v35.0, which served as the domain database^29^.

### Comparison of gene models between our assembly and the Ensembl assembly

miniprot^29^ was ran to map the translated proteins predicted from our assembly onto the female NMR genome in Ensembl. Subsequently, bedtools intersect (v 2.31.1)^30^ with ‘-s -loj’ options was used to identify overlapping regions between the mapped proteins and GFF3 file of the female NMR genome in Ensembl release-111 (accessed on May 30, 2024). Amino acid sequences were aligned using MAFFT^31^ with ‘--auto’ option, and then the alignments were visually examined to evaluate the differences of gene structures between our assembly and the female NMR genome in Ensembl.

### Comparison of genomic differences between our assembly and the Ensembl assembly

Dot plots were generated to visualize the genomic differences between our assembly and the Ensembl assembly. To evaluate genomic differences within the NMR, dot plots were generated among mouse strains (C57BL/6J vs. 129S1/SvilmJ_v1, C57BL/6J vs. A/J_v1, C57BL/6J vs. AKR_J_v1, and C57BL/6J vs. BALB_cJ_v1) and between mouse species (C57BL/6J vs. *Mus spretus* (SPRET/EiJ)). The same process was applied to combinations of mouse strains to evaluate intraspecific differences in NMR. C57BL/6J in Ensembl release-111^34^, GCA_001624185.1 (129S1/SvImJ_v1)^35^, GCA_001624215.1 (A/J_v1)^36^, GCA_001624295.1 (AKR/J_v1)^37^, GCA_001632525.1 (BALB/cJ_v1)^38^, and GCA_921997135.2 (SPRET_EiJ_v3)^39^ were used in this analysis. Minimap2 v2.28^40^ was used to align genome sequences with the ‘-cx asm5’ option. Subsequently, dot plots were visualized using dotPlotly with ‘-s -t -m 500 -q 500000 -l’ options (https://github.com/tpoorten/dotPlotly, accessed on July 17, 2023).

In addition, structural variants (SV) between our assembly and the Ensembl assembly were searched. Minimap2 v2.28^40^ with ‘asm5 --cs -r2k’ option was used to map our assembly to female NMR assembly in Ensembl. Svim-asm v1.0.3^41^, using the default option, was used to identify SVs greater than 40 bp.

### Search for sequence differences of genes between our assembly and the Ensembl assembly

To compare the gene sequences between our genome assembly and the female NMR assembly in Ensembl, the gene annotations of our assembly were transferred to the female NMR assembly in Ensembl using Liftoff v1.5.2^43^. Transferred gene annotations were filtered using the parameter “valid_ORF=True” and “match_ref_proteins=FALSE” to identify transcripts with intraspecific variations of translated amino acid sequence between our assembly and female NMR assembly in Ensembl. The same processes were applied to combinations of mouse strains (C57BL/6J vs. 129S1/SvilmJ_v1, C57BL/6J vs. A/J_v1, C57BL/6J vs. AKR_J_v1, and C57BL/6J vs. BALB_cJ_v1) to evaluate intraspecific differences in NMR. C57BL/6J in Ensembl release-111^34^, GCA_001624185.1 (129S1/SvImJ_v1)^35^, GCA_001624215.1 (A/J_v1)^36^, GCA_001624295.1 (AKR/J_v1)^37^, and GCA_001632525.1 (BALB/cJ_v1)^38^ were used in this analysis. The transcripts with intraspecific variations in NMR were filtered using the Liftoff parameter “Sequence_ID,” which indicates the sequence identity. The transcript ID of NMR and each mouse strain were mapped to those of the C57BL/6J protein ID using the annotation results from Fanflow^44^ and Ensembl Biomart, respectively. Intervene^45^ (accessed October 18, 2024) was used to identify the intersections of the gene sets. Enrichment analysis was performed using the Metascape software with default settings (https://metascape.org/, accessed on October 25, 2024)^46^.

## Results and Discussion

### Sequencing and genome assembly

We obtained 429.39Gb of raw data of short-read sequencing. This data size corresponds to 167.13 times the genome coverage. GenomeScope2^18^ (k=32) estimated the genome size to be 2.569 Gb, with 18% consisting of repeat sequences (Figure 1). The rate of heterozygosity was 0.173%. In addition, a total of 15.2M subreads were generated from two SMRT cells, yielding 282.41 Gb raw data (109.9-fold whole-genome coverage). The final assembly of those subreads resulted in a genome size of 2.561 Gb, which was comparable to that estimated by GenomeScope2^18^ (Table 1). We compared the assembly statistics of the current and conventional assemblies in Ensembl release-111 and release-109 (Table 1). Contiguity of our genome assembly was inferior to that of the current Ensembl genome (release-111), but Benchmarking Universal Single–Copy Orthologs (BUSCO)^22^ showed our assembly had the highest completeness (95.2%).

**Figure 1.**
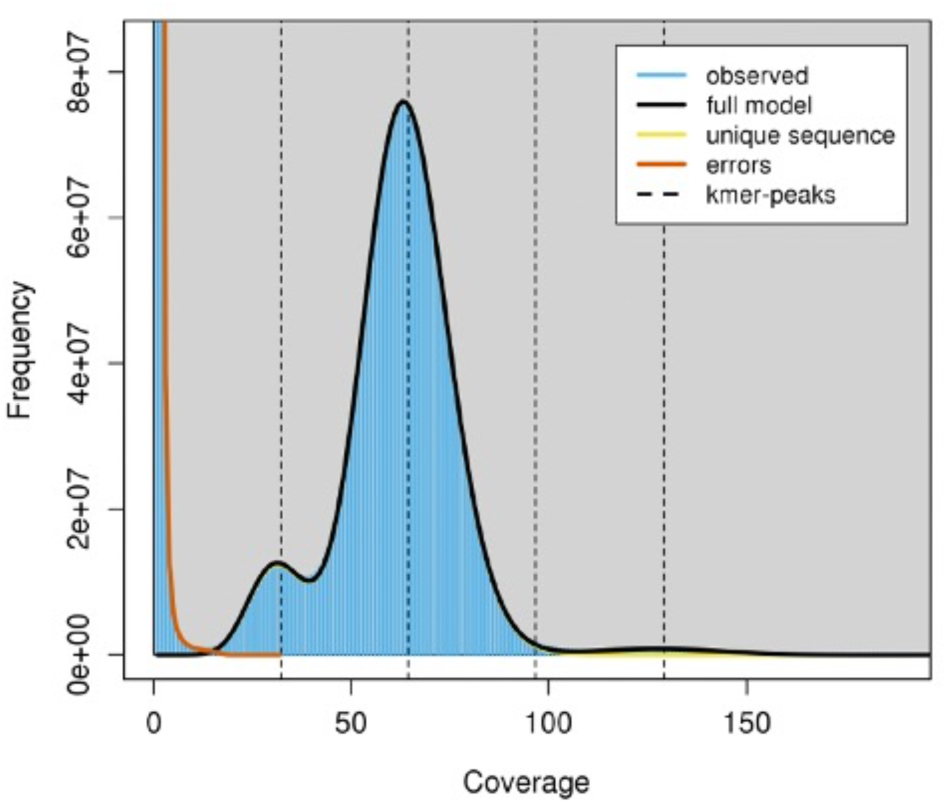
GenomeScope2 results for genome sizes, heterozygosity rate, and repeat sequences Heterozygous and homozygous peaks are shown by the first and second peaks, respectively.

**Table 1.**
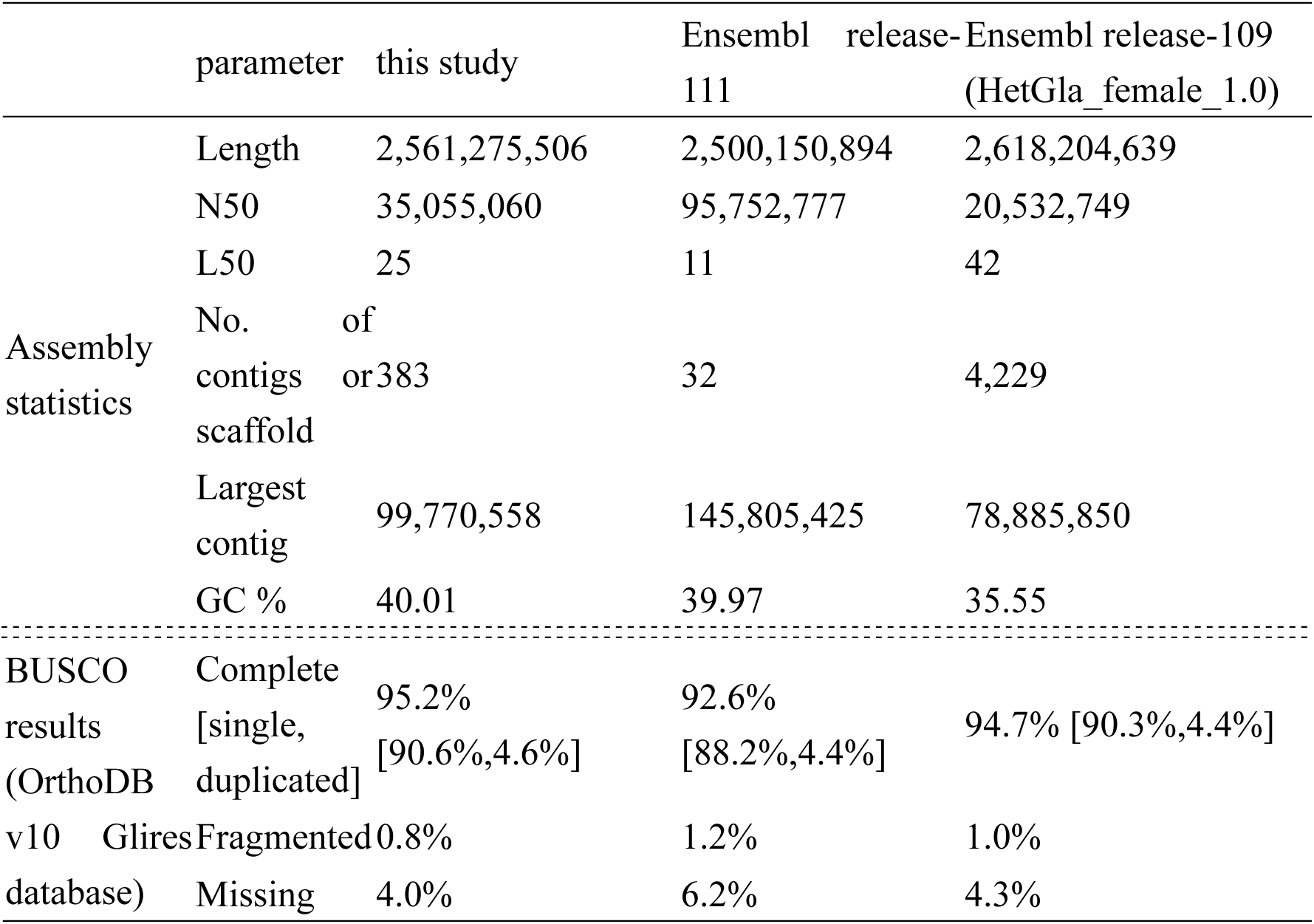
Comparison of assembly statistics with reference genomes in Ensembl

### Gene prediction and functional annotation

BRAKER3 predicted 26,714 protein-coding genes in the assembly (Table 2). BUSCO completeness values were comparable to those of the current and conventional genomes in Ensembl. Of the 26,714 predicted genes, 98% (26,232/26,714) showed matches to at least one protein in the reference protein sets or the Pfam database, leaving only a small number of unclassified proteins (Table 3).

**Table 2.**
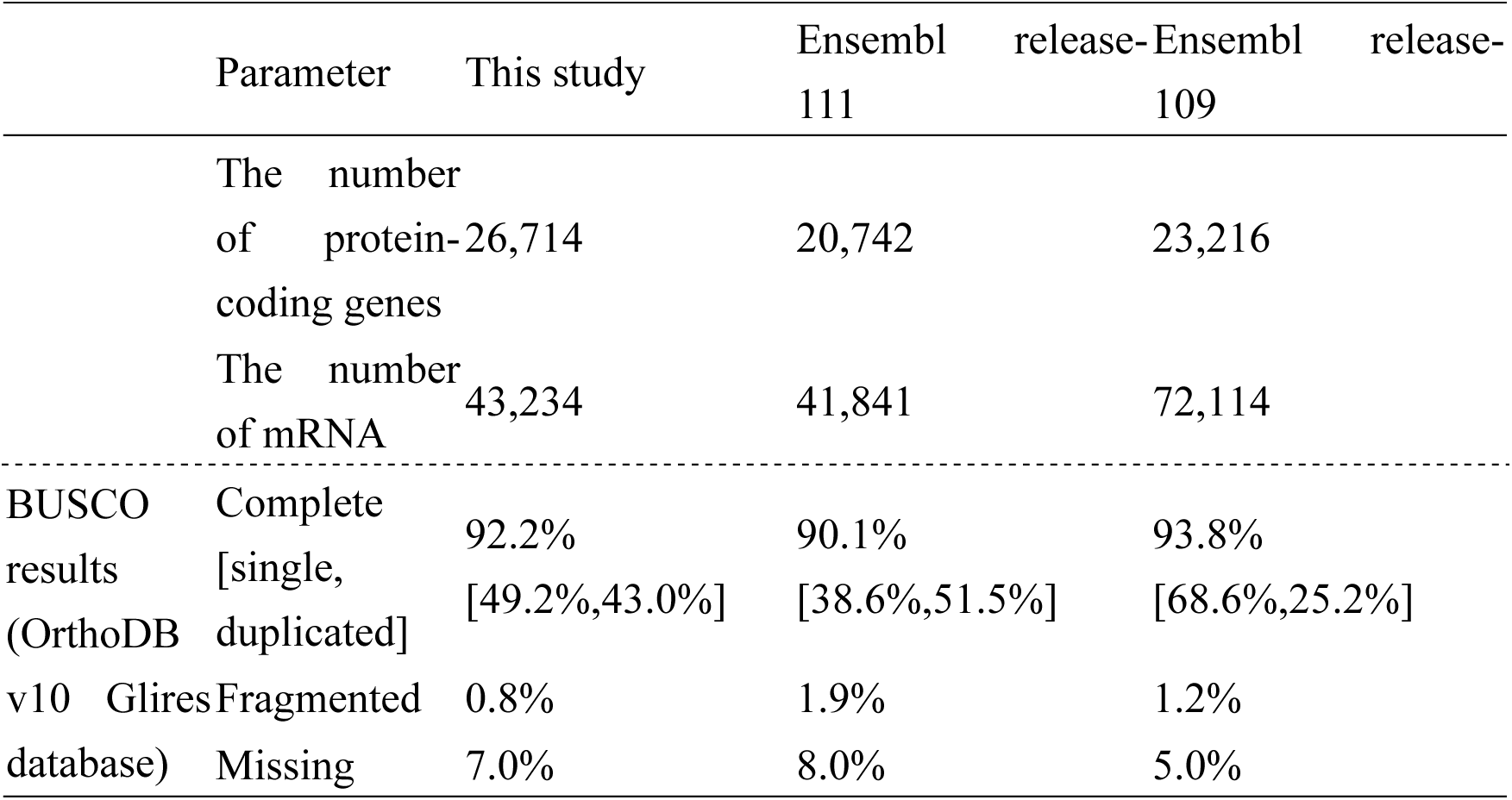
Comparison of gene prediction statistics between our assembly and the reference genome in Ensembl

**Table 3.** Summary of the functional annotation results

| Annotation category | Reference | The number of gene annotated |
| --- | --- | --- |
| 1. Gene homolog (GGSEARCH) | Human | 18,730 |
|  | Mouse | 18,251 |
|  | Guinea pig | 17,080 |
|  | NMR (female) | 17,897 |
|  | NMR (male) | 17,983 |
|  | UniProtKB/Swiss-Prot | 19,149 |
|  | At least one of the above | 26,120 |
| 2. Genes annotated only by HMMSCAN | Pfam domain database | 112 |
| 1 + 2 | - | 26,232 |
| Hypothetical protein | No hit | 482 |
| Total |  | 26,714 |

### Comparison of gene models between our assembly and the Ensembl assembly

During Fanflow execution, 417 transcripts with counterparts in the protein sets of all Ensembl reference species were found, except for those in NMR (female and male). This implied that these transcript models were not present in the Ensembl NMR genome. To confirm whether these 417 transcripts were missing from the NMR genome in Ensembl, we ran miniprot^30^ to map the translated proteins predicted from our assembly onto the female NMR genome in Ensembl. Of the 417 transcripts, 400 were mapped to the female NMR genome in Ensembl, whereas 17 were not (Figure 2a)^33^. The 400 transcripts were further categorized into two groups: one consisting of those present in the gene model of the female NMR genome in Ensembl (250 transcripts), and the other consisting of those absent from the gene models (150 transcripts) (Figure 2a)^33^. Among the 250 transcripts, 39 overlapped across multiple gene models in the female NMR genome of Ensembl (e.g., the g21545 gene, as shown in Figure 2b). The exon of g21545 overlapped with those of ENSHGLG00000049606 and ENSHGLG00000018997. These results show that previously undetected gene models in the Ensembl genome were included in our assembly.

**Figure 2.**
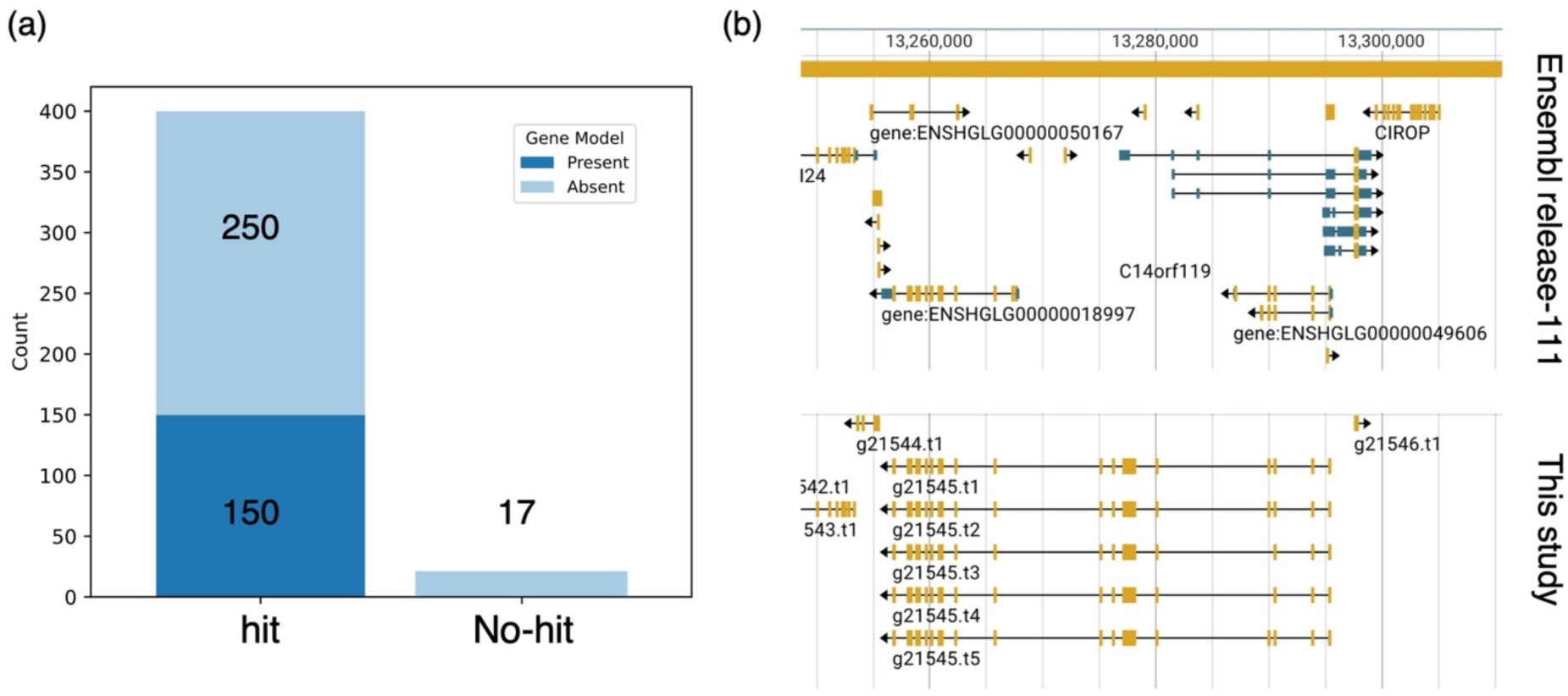
Searches for the gene model differences between our assembly and the Ensembl assembly (release-111) The number of regions mapped by miniprot is shown, with columns colored differently to indicate the presence of overlaps with the Ensembl gene model (a). Gene structures of g21545 and their corresponding regions in the Ensembl genome are shown (b). Yellow and blue boxes indicate the exons and untranslated regions (UTRs), respectively.

### Comparison of genomic differences between our assembly and the Ensembl assembly

Several inversions were observed in the NMR genome, but most plots showed a nearly linear pattern (Figure 3a-c). In contrast, the dot plot between mouse strains and mouse species showed an almost completely linear pattern (Figure 3d-g). These results suggest the presence of intraspecific chromosomal variations within the NMR species. As a result of Svim-asm, 2,263 deletions and 2,668 insertions were identified between our assembly and the Ensembl assembly (Table 4). To evaluate these statistics, we examined the SVs between the mouse strains (Table 4). The number of SVs (deletions or insertions) exceeded 100,000 in all mouse pairs examined in this study. The number of SVs (deletions or insertions) was lower in NMR than in the mouse strains, indicating that structural variations within NMR may be smaller than those within mouse strains. Structural variant NMR data are available in figshare^42^.

**Figure 3.**
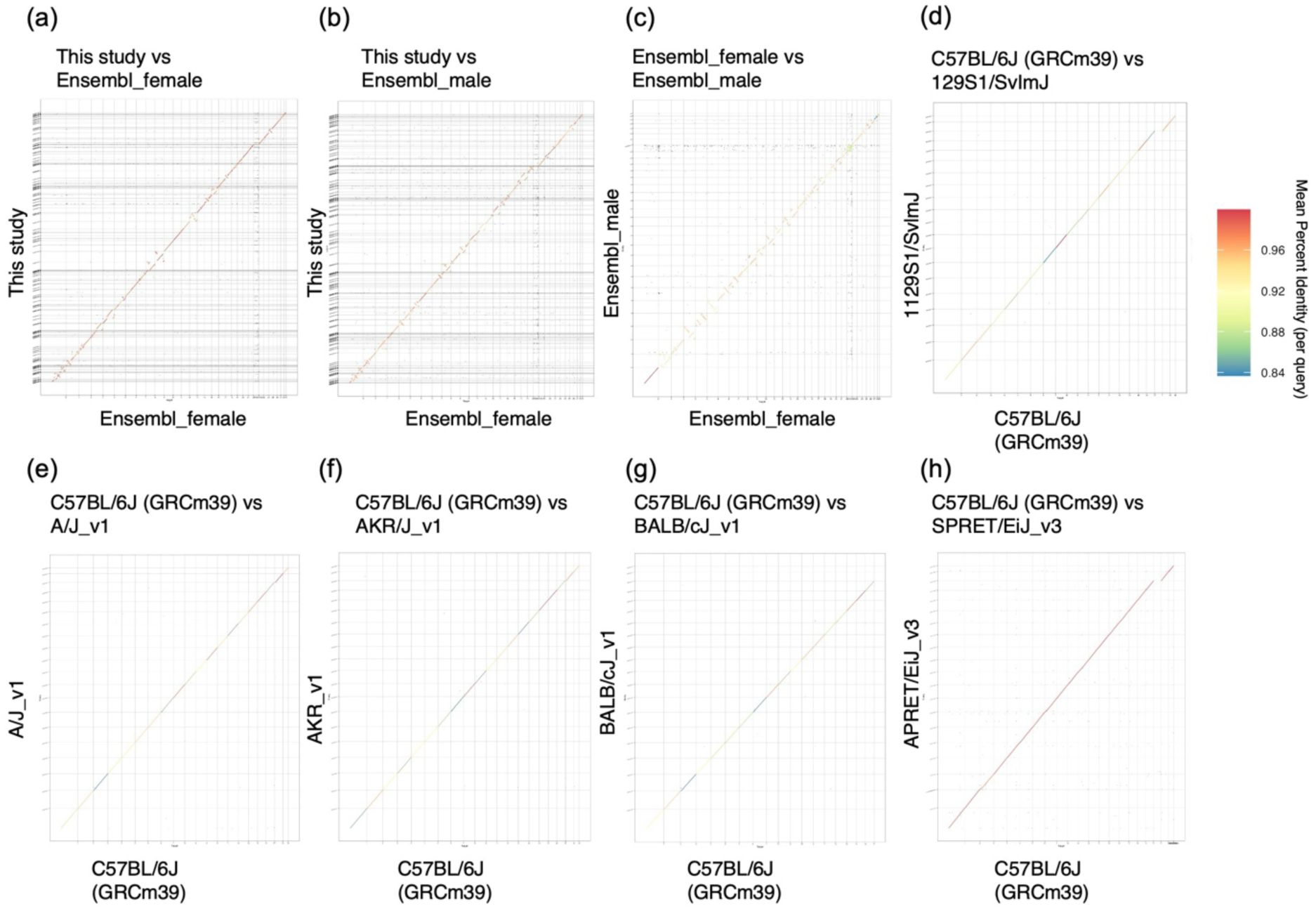
Differences in genome structures within or between species. Dot plots indicate the comparisons of genome structures between our assembly and Ensembl assembly release-111 (female and male) (a-c), between C57BL/6J (GRCm39) and each mouse strain (d-g), and between C57BL/6J (GRCm39) and SPRET/EiJ (*Mus spretus*) (h). The color of plots indicates the mean percent identity (per query).

**Table 4.**
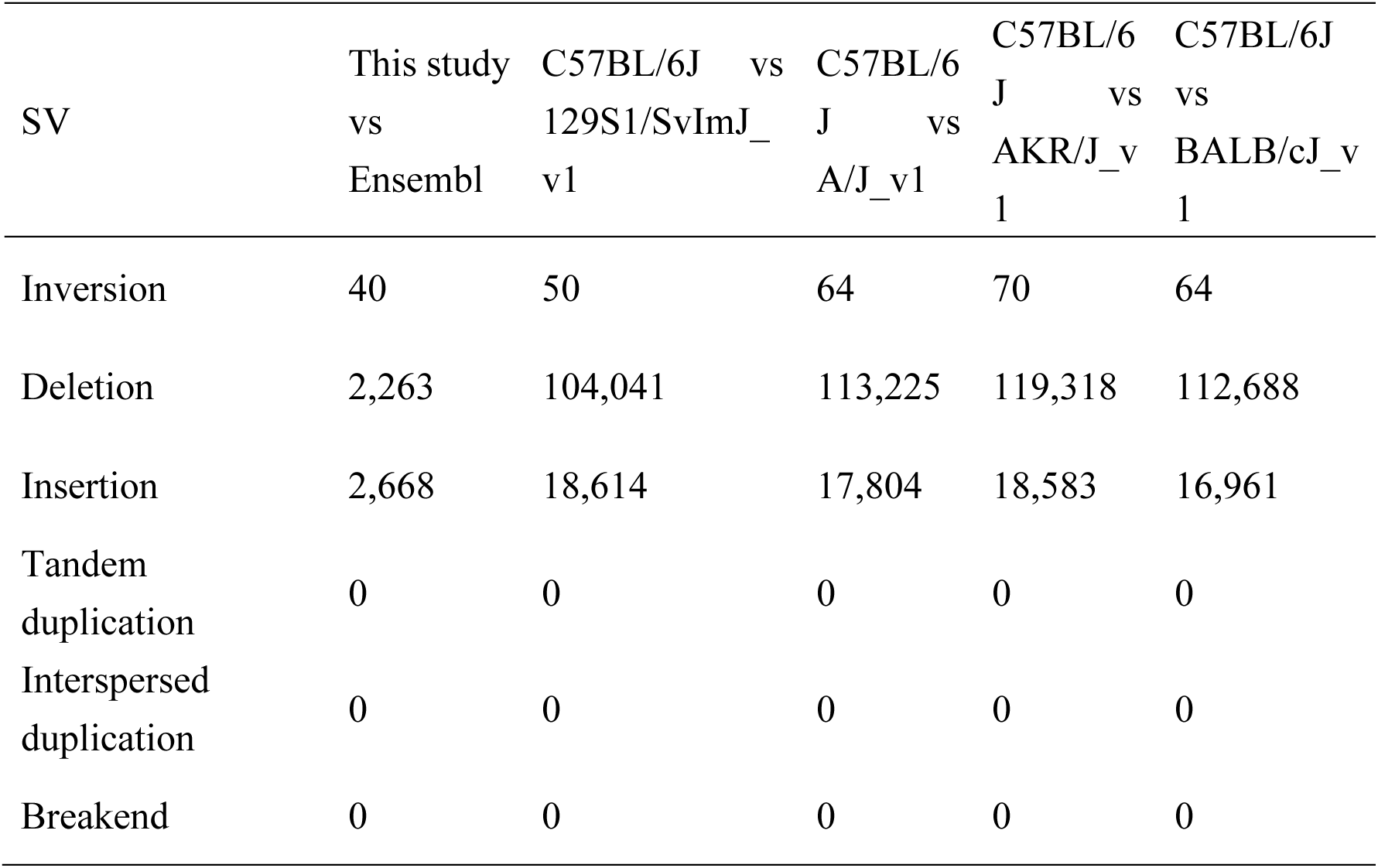
Structural variants within NMR and within mice

### Search for sequence differences of genes between our assembly and the Ensembl assembly

The annotation of 32,263 transcripts from our assembly was successfully transferred to the female NMR assembly in Ensembl, and sorted by indicators of sequence identity, “Sequence_ID” (Figure 4a). In the NMR comparison, 177 transcripts had a “Sequence_ID” below 0.983, and the count per “Sequence_ID” below this threshold was consistently under 10, suggesting that intraspecific variation in these transcripts was pronounced in the gene sets analyzed in this study. When comparing intraspecific transcripts between NMR and mouse strains, 77 genes exhibited NMR-specific intraspecific variations (Figure 4b)^47^. Enrichment analysis of the 77 transcripts revealed significant enrichment of genes involved in amine ligand-binding receptors, including 5-hydroxytryptamine (serotonin) receptor 2B (*Htr2b*), G protein-coupled receptor 143 (*Gpr143*, a receptor for L-3,4-dihydroxyphenylalanine [L-DOPA]), erythropoietin (*Epo*), and thyroid hormone receptor beta (*Thrb*) (Figure 4c)^48^. Intraspecific variations in *Htr2b* and *Gpr143* suggest that responses to serotonin and L-DOPA may differ even within NMR populations, potentially influencing social behavior. Notably, NMR fibroblasts exhibit unique serotonin management, in which serotonin accumulates and induces cell death through serotonin metabolism in response to cellular senescence^49^. Additionally, intraspecific variations in *Epo* and *Thrb* may contribute to adaptation to hypoxic underground environments through changes in erythropoiesis and metabolic regulation. Examining the functions of these genes could help deepen our understanding of the molecular mechanisms underlying the unique traits of NMRs, such as eusociality, hypoxia tolerance, and delayed aging. Overall, genes exhibiting intraspecific variations may provide valuable insights into the behavioral and physiological adaptations of NMR.

**Figure 4.**
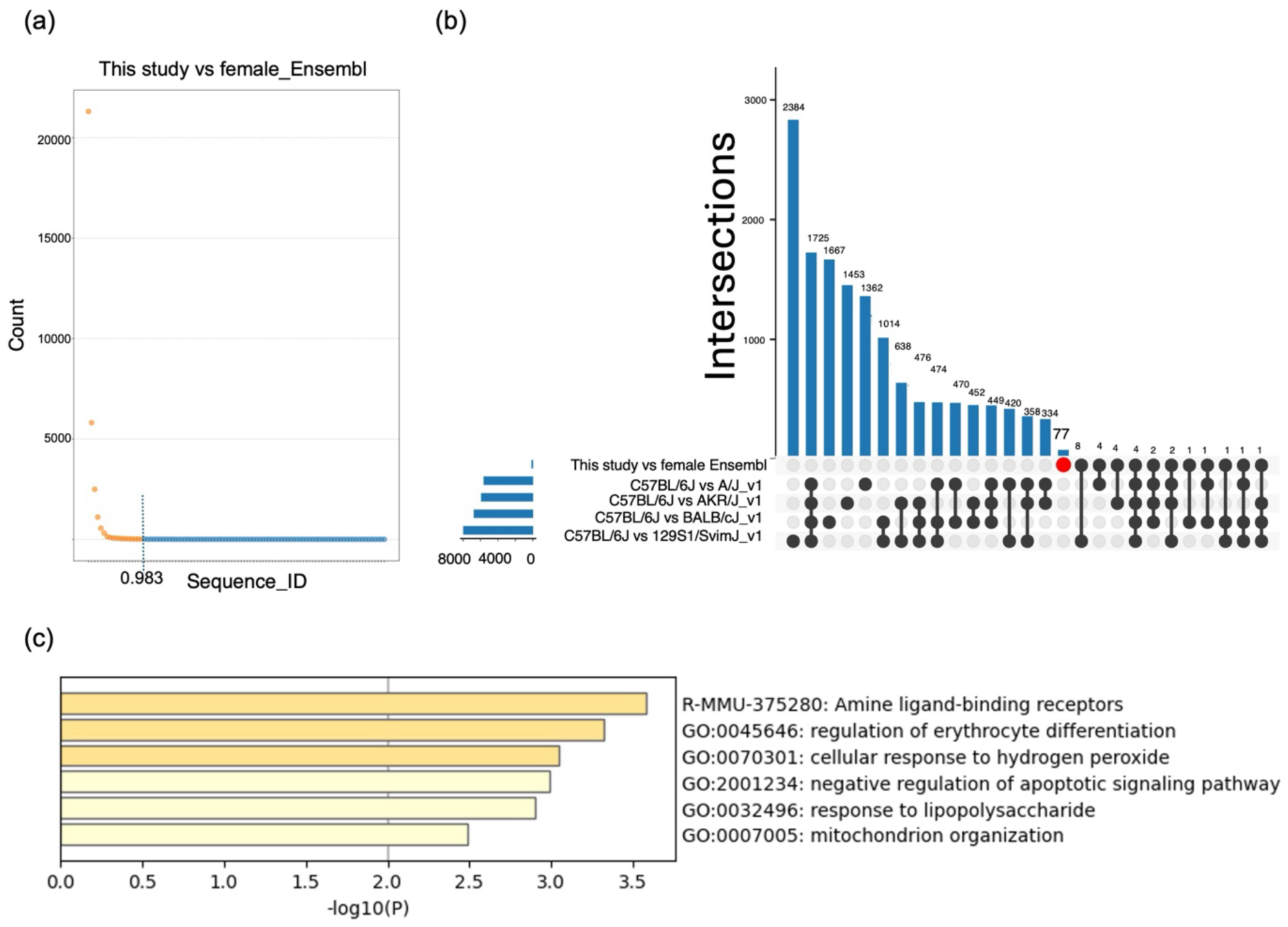
Mutations for amino acid sequences between our assembly and the female Ensembl assembly. (a) Total numbers of transcripts per Sequence_ID. The dot color changes based on a Sequence_ID threshold of 0.983. (b) Intersections of transcripts with intraspecific variations in NMR and mouse strains. Red circle indicates transcript sets with intraspecific differences only within NMR. (c) Enriched terms in NMR-intraspecific 77 transcripts indicated by red circle in (b).

## Data Records

All raw sequence data for genome assembly (Illumina short reads and PacBio CLR) have been deposited in the DNA Data Bank of Japan under the BioProject accession number PRJDB14267. Full information on functional annotations added using Fanflow is available at *figshare*^44^. Structural variants within NMRs have been deposited in *figshare*^42^. Gene models estimated by BRAKER3 for our assembly have been deposited in *figshare*^50^.

## Technical Validation

The performance of the assemblers was evaluated by comparing four assemblers. The four assemblers (wtdbg2^51^, Flye v2.8^52^, Canu v2.0^19^, and Canu v2.1^19^) were used to compare the assembly quality with respect to contiguity. The assembly, constructed by Canu v2.1, had the longest average and maximum lengths in contigs, and showed the minimum L50 value (Table 5). Pilon was used to polish the genome assembly constructed using Canu v2.1, and Purge_dups^21^ was used to remove haplotigs. This process improved genome statistics, particularly regarding contiguity (Table 6).

**Table 5.**
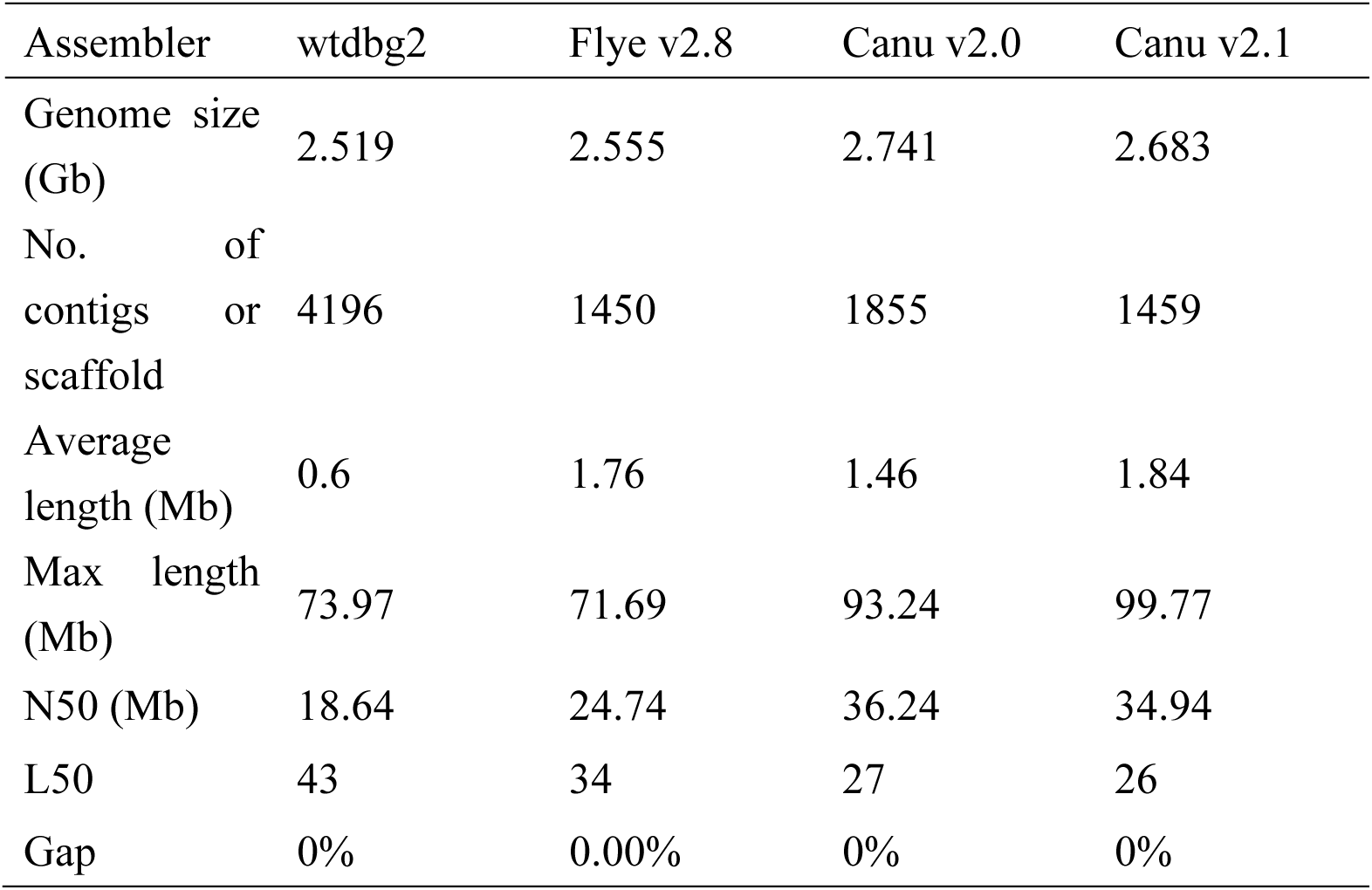
Comparison of genome assemblers

**Table 6.** Results of polishing using short reads and removal of haplotigs with Purge_dups

|  | Before | Pilon (three times) | Purge dups |
| --- | --- | --- | --- |
| Genome size (Gb) | 2.683 | 2.683 | 2.561 |
| No. of contigs or scaffold | 1459 | 1459 | 383 |
| Average length (Mb) | 1.84 | 1.84 | 6.69 |
| Max length (Mb) | 99.77 | 99.77 | 99.77 |
| N50(Mb) | 34.94 | 34.94 | 35.06 |
| L50 | 26 | 26 | 25 |
| Gap (%) | 0 | 0 | 0.0000012 |

## Code availability

No specific programs or codes were used in this study.

## Acknowledgments

This work was supported by JSPS KAKENHI Grant Number JP16H06279(PAGS), supported in part by the JSPS KAKENHI Grants (JP18H02365) to K.M., JSPS KAKENHI Grants (JP24H00542) and JST COI-NEXT (JPMJPF2010) to K.M. and H.B. and JST FOREST Program (JPMJFR216C) to K.M. We would like to thank all laboratory members at Hiroshima University and Kumamoto University for their valuable comments. Computations were performed on the computers at the Hiroshima University Genome Editing Innovation Center. Computations were partially performed on the NIG supercomputer at ROIS National Institute of Genetics.

## Author Contributions

K.M. and H.B. coordinated and designed this study. K.O. conducted the sampling of *Heterocephalus glaber* and H.T., T.I., and A.T. performed sequencing and de novo assembly. K.T. and H.B. performed bioinformatics analyses and generated figures and tables. K.T. wrote the first draft of the manuscript. All authors revised, edited, and approved the final manuscript.

